# Suppressing selection for antibiotic resistance in the environment: A transparent, ecology-based approach to predicted no-effect concentrations

**DOI:** 10.1101/2025.04.04.647007

**Authors:** David Kneis, Magali de la Cruz Barron, Diala Konyali, Valentin Westphal, Patrick Schröder, Kathi Westphal-Settele, Jens Schönfeld, Dirk Jungmann, Thomas U. Berendonk, Uli Klümper

**Affiliations:** Technische Universität Dresden, Institute of Hydrobiology, Dresden, Zellescher Weg 40, Germany; German Environmental Agency (UBA), Section IV2.2 Arzneimittel, Wörlitzer Platz 1, 06844 Dessau-Roßlau, Germany

**Keywords:** Antibiotic resistance, Environmental risk assessment, Minimal selective concentration, Predicted no effect concentration, Resistance cost, Environmental Regulation

## Abstract

Selection for antibiotic resistance has been demonstrated at low, environmentally-relevant antibiotic concentrations. Over the past decade, the concept of minimum selective concentrations (MSC) has been adopted in environmental regulation to define maximum permissible antibiotic concentrations. Such empirically determined MSC values often fail to reflect the complexity of natural communities, where susceptibility and resistance-associated fitness costs vary widely across species. To address this limitation, computational approaches have been developed to predict no-effect concentrations for selection of antibiotic resistance (PNEC_res_) from routinely collected minimum inhibitory concentration (MIC) data. However, these approaches often lack a strong ecological basis, undermining confidence in their predictions.

Here, we propose a simple but biologically consistent framework to derive PNEC_res_ values by integrating MIC data with probabilistic estimates of resistance-related fitness costs. Our results suggest that current regulatory environmental threshold concentrations should be lowered by at least one order of magnitude to guard against selection for antibiotic resistance.

## Introduction

The widespread occurrence of antimicrobial resistance (AMR) across human and animal microbial pathogens is currently leading to a global health crisis^1^. In 2021 alone, around 4.7 million human deaths were associated with bacterial AMR, with 1.1 million of these directly attributable to bacterial AMR, with numbers predicted to increase to 1.9 million deaths attributable and 8.2 million deaths associated with AMR globally in 2050^2,3^. Within recent years, it has become clear that the ongoing spread of AMR is not only associated with the clinical or veterinary sphere but also happens within and through natural environments^4–7^. Within these environments, it is crucial to note that, unlike chemical pollutants, antimicrobial resistance genes (ARGs) are potentially amplified through bacterial growth and lateral gene transfer^6,7^. This makes defining safe concentrations of antibiotics in the environment particularly challenging.

Positive selection for bacteria hosting ARGs occurs already at very low, sub-inhibitory concentrations of antibiotics^8,9^. Antibiotic concentrations high enough to select for AMR can be found in numerous matrices entering the environment, such as wastewater from antibiotic production sites, effluent from municipal wastewater treatment plants or manure^10,11^. This makes the environment not only a conduit for antibiotic-resistant bacteria (ARB) and their ARGs introduced through anthropogenic sources but also a reactor for their enrichment in environmental microbiomes from which they can thereafter return to the human or animal spheres.

Consequently, to limit selection for AMR in the environment there is an urgent need for quantitative risk-based assessment and subsequent policy-based regulation of antibiotic pollution levels^7^. In principle, the need for addressing this problem has already been recognised by legislation. For example, the European Union (EU) authorisation procedure for new veterinary antimicrobial medicinal products already states that “Resistance in the environment shall be addressed”^12^. Also, in the new draft EU legislation on medicinal products for human use, assessing the risks of these products regarding their contribution to the spread of AMR “during manufacturing, use and disposal” shall be included^13^. To date, however, there is no standardised method for carrying out such a risk assessment regarding which concentrations cause selection for AMR. Consequently, there is currently a lack of risk mitigation measures or other regulatory consequences for the application of antibiotics that could reduce the risk for the further spread of AMR via the environment.

To allow the implementation of regulatory thresholds for the environment, establishing predicted no-effect concentrations regarding positive selection of resistance (PNEC_res_) for essentially all antibiotics is paramount. Ideally, PNEC_res_ would accurately display the highest possible concentration of an antibiotic in which no positive selection for any ARG providing resistance against this specific antibiotic occurs across bacterial strains in the environment. However, determining such concentrations for all combinations among the billions of bacterial species, hundreds of antibiotics, and thousands of different ARGs is obviously illusive. Consequently, values of PNEC_res_ are currently derived in a pragmatic manner either from scarce experimental data or by computational prediction. The two alternatives have recently been reviewed extensively and critically by Murray et al.^14^.

Experimental approaches regularly rely on determining empirical minimal selective concentrations (MSCs) based on either competition assays^8,9,15^ or the measurement of differences in growth rates^16,17^ across a gradient of antibiotic exposure. The pairs of strains employed in such competition experiments are generally isogenic except for a chromosomal or plasmid-borne resistance determinant. Approaches aiming for a higher level of environmental realism performed similar competition experiments between isogenic strains in the context of a complex background microbial community^18–21^, resulting regularly in higher MSCs compared to single-strain experiments due to elevated costs of being resistant in competition with other community members. This implies that empirical MSCs determined from single-strain experiments are likely conservative, which could be a benefit from the regulatory perspective when utilizing them to determine PNEC_res_. Approaches based on isogenic strains are highly accurate in empirically determining individual MSCs, but they are also very labour-intensive. Moreover, they will never be exhaustive as they exclusively reveal the MSC for a single strain-ARG combination, bringing into question how transferable results from a single strain are to the extremes observed across complex environmental microbiomes.

Other empirical approaches rely on experimentally evolving environmental microbial communities across a gradient of antibiotic concentrations and either directly determining the MSC based on dynamic changes in the relative abundance of ARGs within the communities^22–24^ or indirectly estimating MSCs based on the growth inhibition patterns of the exposed communities^25,26^. However, MSCs derived from these methods are highly dependent on the origin of the microbial community and provide rather a community-wide average of MSCs in the community than display the MSC extremes (e.g., the lowest MSC within the community) that regulatory interventions aim to target.

Computational estimations, on the other hand, can provide an exhaustive picture of PNEC_res_ for a wide variety of different antibiotics. They are moreover comparatively cost-effective, and their prediction based on existing datasets that were collected according to standardized guidelines, such as in the European Committee on Antimicrobial Susceptibility Testing (EUCAST)^27^ or Clinical and Laboratory Standards Institute (CLSI)^28^ databases, improves comparability of the gained PNEC_res_ estimates^14^.

The most widely applied approach to derive estimates of PNEC_res_ was developed by Bengtsson-Palme & Larsson in 2016^29^ and has already been partially adopted for regulatory purposes by the AMR Industry Alliance in their recommendations for threshold antibiotic concentrations in the receiving water of pharmaceutical manufacturing effluent^30^. This approach uses minimal inhibitory concentration (MIC) values extracted from the EUCAST database^27^ to calculate the final PNEC_res_ for resistance selection by dividing the lowest observed MIC by an assessment factor of 10 to obtain a lower-bound estimate of the MSC. The resulting PNEC_res_ estimates are regularly lower, and therefore more conservative, than MSCs originating from experimental approaches^14^, which is mainly due to the good representation of within and across-species variance of MICs by the EUCAST database. By comparison, experimental approaches are far less exhaustive and tend to measure MSCs close to the median of the MSC distribution, whereas minimum values remain unidentified.

A similar method for computationally deriving PNEC_res_ was proposed by Rico et al.^31^. However, they do not exclusively use the lowest reported MIC from the EUCAST database as in Bengtsson-Palme & Larsson^29^. Instead, they first pool the reported MICs at genus level before dividing by factor 10 to obtain an empirical distribution of MSCs^31^. They then apply a species sensitivity distribution to derive PNEC_res_ for individual antibiotics. Specifically, they approximated the log-transformed MSCs of all genera by a normal distribution and defined PNEC_res_ as the theoretical 5% quantile in analogy to the estimation of hazardous concentration (HC_5_). Like with the Bengtsson-Palme & Larsson^29^ approach, the employed assessment factor of 10 to convert MICs into MSCs is lacking a strong biological justification. Moreover, the assumption of normality could not be verified in a substantial number of cases according to the supplementary information of Rico et al.^31^.

The European Food Safety Authority (EFSA) to establish PNEC_res_ for antibiotics in animal feed^32^ is equally based on using the MIC distribution. Rather than adding a plain assessment factor like in the previous two approaches, they apply known ratios of MSC/MIC for experimentally tested species^32^. This makes the derived values of PNEC_res_ in theory more empirically based, but also results in high experimental effort for determining MSC/MIC ratios and brings into question if these ratios determined for the tested bacterial species (often laboratory strains) are indeed transferable to environmental bacterial species.

The computational approaches to the derivation of PNEC_res_ are, in general, more easily applied and more exhaustive than experimental approaches rooted in competition experiments. However, the former still fail to explicitly take into account fundamental evolutionary and ecological mechanisms underlying the spread of ARGs, such as selection, dose-response patterns, and the costs of maintenance of different resistance determinants. Instead, they adopt the concept of assessment factors from classical ecotoxicology and thus employ a universal constant to translate MICs into MSCs. This has led Murray et al.^14^ to conclude in their recent meta-analysis that, until such shortcomings are ruled out, a general PNEC_res_ of 0.01 µg/L for all antibiotics could be considered as a reasonable alternative to the values obtained by any experimental or computational approach.

A first attempt at predicting biologically meaningful MSC/PNEC_res_ estimates was made by Greenfield et al.^33^ who formulated general ecological relationships between MIC, MSC, and the cost associated with resistance. Owing to a considerable number of unknown parameters, however, their approach has not been adopted by regulatory practice so far.

In this paper, we demonstrate with a minimum of basic math that, for typical high-level resistances, the ratio MSC/MIC is generally almost identical to the cost of the resistance determinant. Consequently, biologically meaningful estimates of the MSC can be obtained from simple standard microbiological assays without the need for expensive and time-consuming competition experiments. We provide experimental evidence for the theory-borne relation between MIC and MSC with the example of several different host strains and a set of multiple plasmid-borne and chromosomal resistances of varying costs. Subsequently, based on the statistical analysis of reported resistance costs, we derive a biologically meaningful substitute for the commonly used assessment factor of 10^29–31^. Finally, we discuss remaining challenges toward the establishment of PNEC_res_ for all antibiotics based on evolutionary and ecological processes. Our work opens a road toward a feasible computational derivation of PNEC_res,_ which are consistent with biological principles and can easily be used to calculate different levels of protection. It is thus a significant contribution to future risk assessment and subsequent environmental regulation of antibiotics.

## Results & Discussion

### Expected relationship between MSC and MIC

The MSC is defined as the antibiotic concentration where a susceptible and an antibiotic-resistant bacterial strain are subject to equal fitness represented by identical growth rates (Fig. 1).

**Figure 1:**
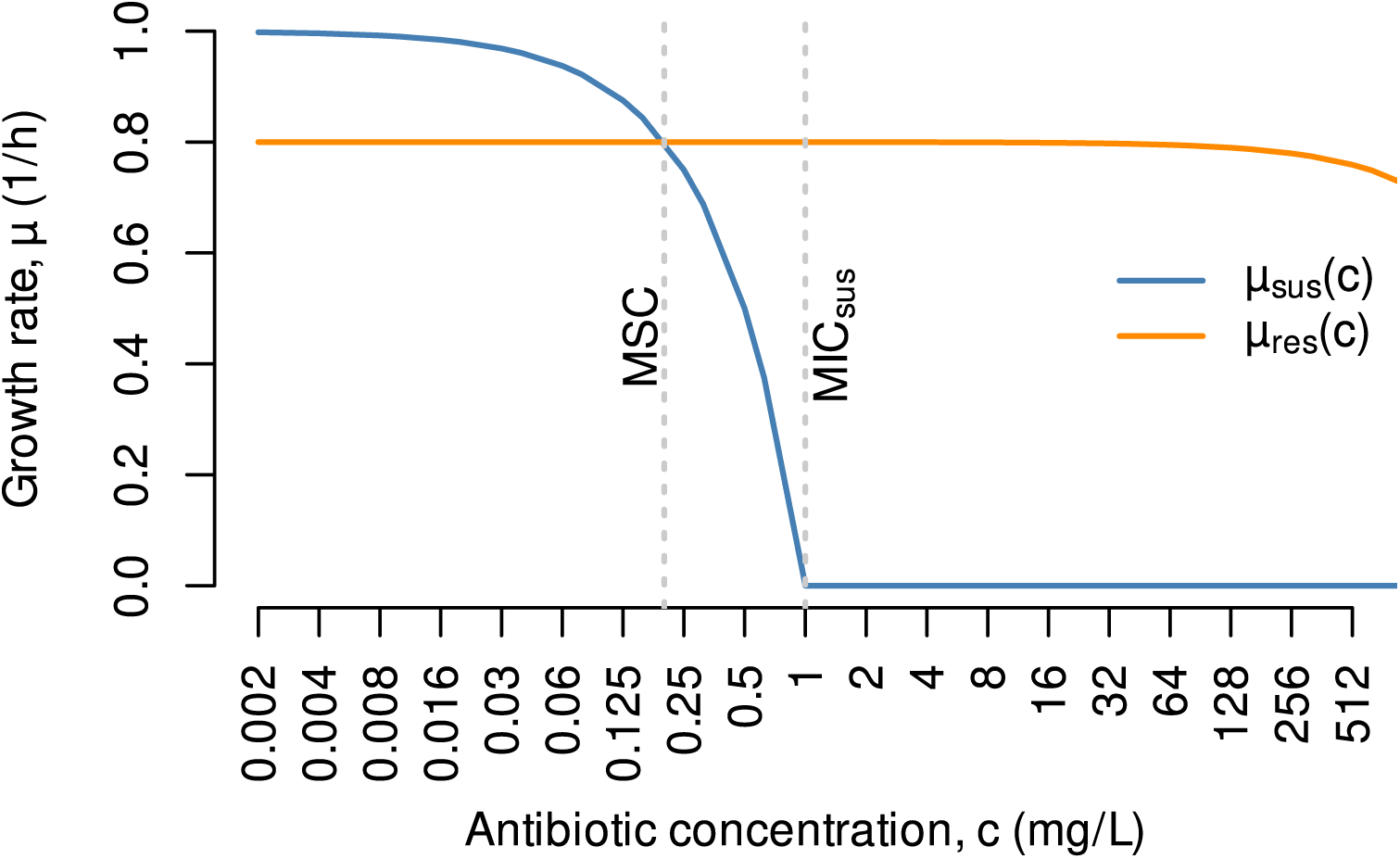
Growth rates (µ) of a susceptible and a resistant strain (subscripts "sus" and "res") across a gradient of antibiotic concentrations (c). The intersection between the two graphs defines the MSC. MIC_sus_ marks the lowest concentration fully inhibiting the growth of the susceptible strain. Note the logarithmic lower axis. Inhibition of the resistant strain is assumed to occur at very high concentrations only (MIC_res_ beyond lower axis range).

The effects of antibiotics on the growth of bacterial strains can regularly be described through simple dose-response models^33^. The most basic one (Fig. 1) assumes a linear relation between growth rates (µ; unit: h^-1^) and the concentration of the antibiotic (c) as expressed by Eq. 1. The linear model has two parameters: the intrinsic maximum growth rate at zero antibiotic exposure (µ^0^) and the minimum inhibitory concentration (MIC).

The shape of actual dose-response curves differs between antibiotics, and it is potentially influenced by various other factors, including phenotypic characteristics and general growth conditions. Besides linear dose-response relations, there are cases where a measurable decline in growth only emerges at concentrations close to the MIC and, in yet other cases, even very low doses of antibiotics trigger a notable drop in fitness^34^ (Fig. S1). In the following, we focus on the linear dose-response relation (Eq. 1) as a parsimonious consensus model and discuss its applicability. Since antibiotic concentrations beyond the MIC are hardly relevant in environmental settings, the values of µ obtained by Eq. 1 are generally positive.

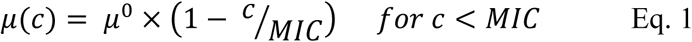

Let the subscript "res" denote an antibiotic-resistant and the subscript "sus" a susceptible variant of an otherwise isogenic pair of bacterial strains associated with individual intrinsic growth rates (µ^0^_res_, µ^0^_sus_) and minimum inhibitory concentrations (MIC_res_, MIC_sus_). Further, let µ^0^_res_ be related to µ^0^_sus_ by a dimensionless cost factor in the range 0-1 representing the total fitness burden associated with the resistance determinant, be it a single ARG or an AMR encoding plasmid (Eq. 2). For example, a cost factor of 0.05 would imply a 5% fitness burden of the resistance determinant on the strains growth rate.

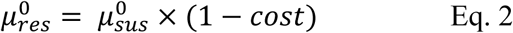

The definition of the minimum selective concentration (MSC) implies the equality in growth rates between the resistant and the susceptible strain expressed by Eq. 3 (Fig. 1). Making use of Eq. 1 and Eq. 2, both sides of Eq. 3 can be expanded to yield Eq. 4 relating the MSC to measurable quantities.

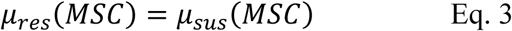

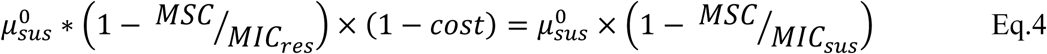

To simplify Eq. 4, we introduce a positive dimensionless factor f, representing the ratio of the two strains’ MICs (Eq. 5). Specifically, f expresses the level of antibiotic resistance such that, for example, a value of f=10 indicates a ten-fold increase of the MIC upon acquisition of the particular resistance determinant.

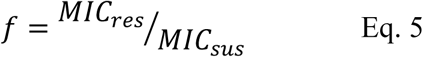

Inserting Eq. 5 into Eq. 4 allows the MSC to be expressed as a function of two parameters: the cost of resistance (Eq. 2), and the level of resistance (factor f, Eq. 5). A particularly handy expression is obtained if the MSC is expressed in normalized form as the quotient MSC/MIC_sus_ (Eq. 6). This displays by which factor the MSC is lower than the MIC of the susceptible strain, which in previous experimental studies was highly variable ranging from as low as 4-fold to several 100-fold lower^8,9,15,18,23,35^.

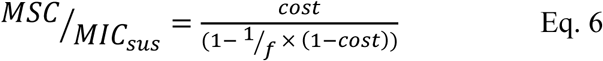

A formal analysis of Eq. 6 reveals that, for large values of factor f, the denominator on the right-hand side approaches unity. Consequently, if the level of antibiotic resistance falls within the range of f > 10, representing more than 94% of the typically observed high-level resistance determinants of clinical relevance^36^, the normalized MSC is numerically very close to the cost of resistance (Fig. 2) giving rise to further simplification (Eq. 7). Note that Eq. 7 can also be derived from Eq. 1 and Eq. 2 by plain geometric considerations known as the intercept theorem.

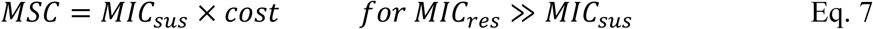

**Figure 2:**
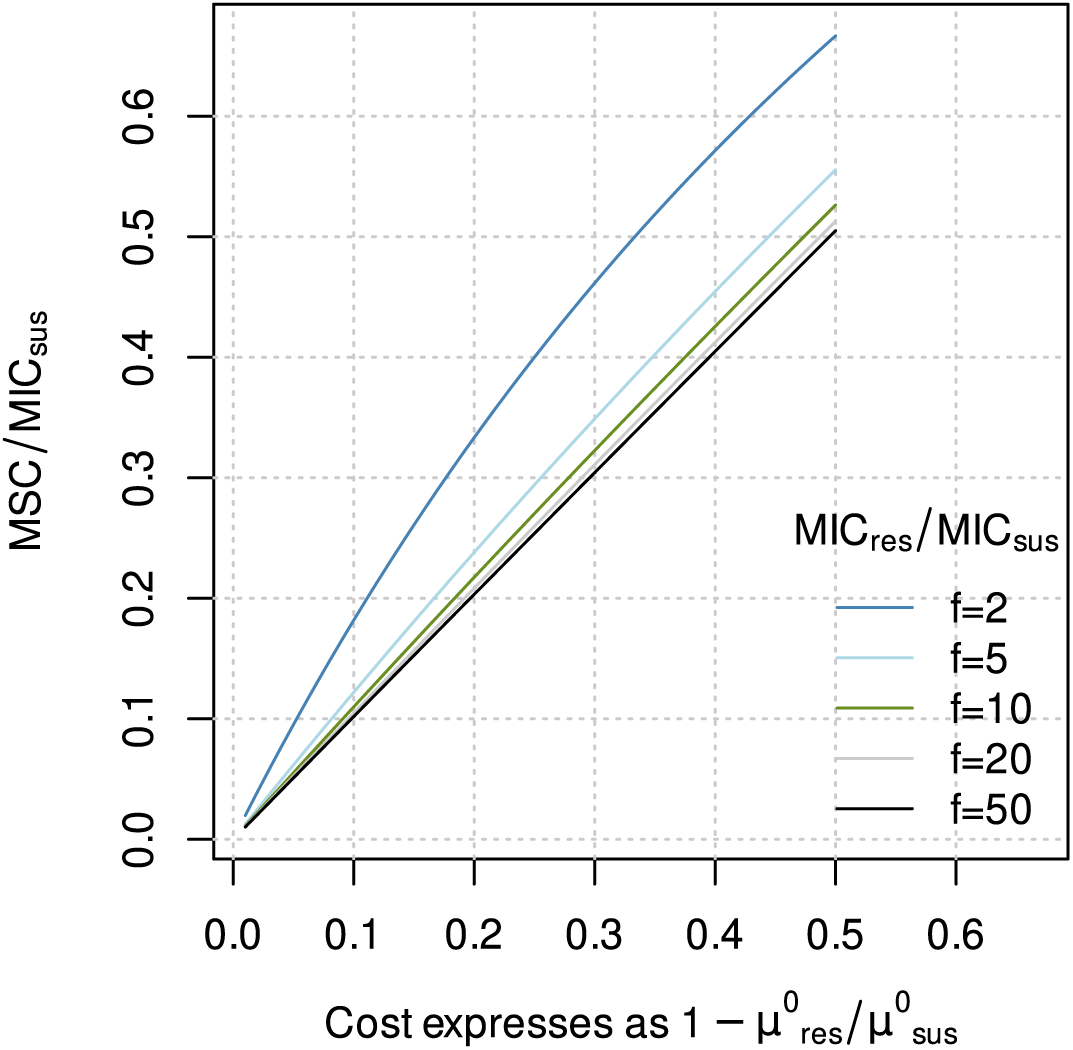
Graphical representation of Eq. 6 illustrating the dependency of the normalized MSC on the cost of resistance (lower axis) and the level of resistance (f) represented by the ratio MIC_res_ /MIC_sus_ (individual curves; cf. Eq. 5).

The practical benefit of Eq. 7 lies in the fact that the MSC can be simply calculated from standard growth assays and MIC data, thus avoiding time-consuming and expensive empirical determination through competition experiments.

### Validation of computed MSCs against empirical MSCs

To verify the practicality of Eq. 7, we compared empirically determined MSCs from classical competition experiments across an antibiotic gradient to their computed counterparts. Computed MSCs were according to Eq. 7 calculated exclusively based on the MICs and the cost of resistance obtained from the same competition experiments. The latter covers 26 strain-antibiotic combinations made up of three model focal bacterial strains (*Escherichia coli*, *Pseudomonas putida*, *Bacillus subtilis*) and resistances to 13 different antibiotics representing six classes of antibiotic drugs encoded through ARGs that were introduced either through plasmids or transposition into the chromosome (see Methods). In 2/3 of the tested cases, the deviation between the computed and empirical MSC was less than factor two (Fig. 3). Thus, the deviation remained below the resolution of the scale commonly employed in the EUCAST^27^ or the CLSI^28^ databases, where MICs are exclusively reported using standard two-fold dilutions of antibiotic concentrations. Deviations exceeding factor four were encountered rarely (12% of the cases). Overall, we determined a low mean error of 0.13 orders of magnitude and a mean absolute error of 0.27 orders of magnitude of the empirical MSCs compared to the computed MSCs. Hence, the empirical results provide reasonable support for the proposed equality of the MSC/MIC ratio and the costs of resistance (Eq. 7).

**Figure 3:**
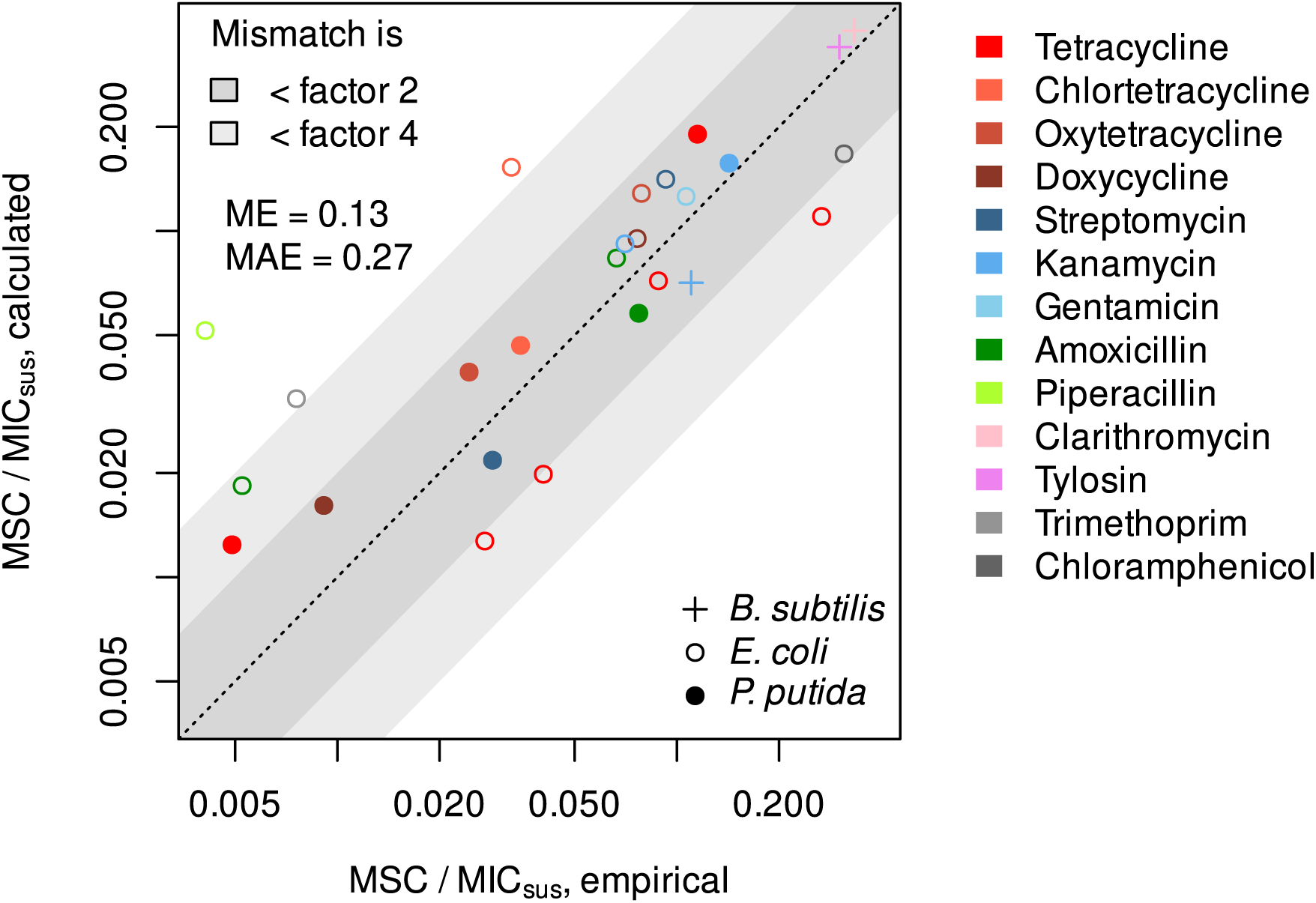
Minimum selective concentrations measured empirically by competition experiments in comparison to values computed by Eq. 7. Mean error (ME) and mean absolute error (MAE) were computed from individual mismatches expressed as log_10_(computed/empirical) to accommodate the wide data range.

### From computed MSCs to PNEC_res_

As outlined above, the MSC represents the critical antibiotic concentration above which selection for resistance occurs. In general, the MSC pertains to a particular pair of bacterial strains that differ concerning the resistance trait but are otherwise isogenic. Nevertheless, it seems attractive to exploit the MSC concept to derive PNEC_res_ of antibiotics in environmental systems targeted at minimizing the likelihood of undesired selection for resistance.

Soils, water, and even wastewater, however, host a large number of bacterial species that differ substantially in terms of antibiotic susceptibility. Crucially, pronounced variation in MICs is not only observed between species but also within a species, as illustrated by the spread of reported MIC distributions in the EUCAST database^27^ and is likely even larger than illustrated there due to the limited representation of environmental isolates in the database. An even larger variability is anticipated for the cost of resistance, which is determined by, e.g., the resistance mechanism, the genetic context of the resistance gene, the characteristic of particular alleles, and finally the bacterial host^37,38^. Moreover, the cost of resistance in a strain can be ameliorated through evolution either rapidly or over time, especially when encoded on plasmids^39,40^. Considering precautionary principles, a PNEC_res_ should thus be based on the lowest MSC among bacteria present in real-world environmental communities.

Given that the phenotypic properties of a substantial fraction of environmental bacteria are effectively inaccessible, we thus propose to derive a PNEC_res_ for a particular antibiotic from the lowest known MIC, thereafter denoted MIC_lowest_, and a lower-bound estimate of the cost of resistance (Eq. 8). For the latter, we propose to adopt a quantile Q_p_ of the overall distribution of resistance costs associated with a sensible probability threshold p (e.g., p = 0.05). Note that Eq. 8 is structurally identical to the approach of Bengtsson-Palme & Larsson, and the major advancement is solely in the rationale behind the factor MIC_lowest_ is adjusted with. In their case, it is exclusively a fixed assessment factor of 10 to account for differences between MICs and MSCs^29^. Here, it represents an ecologically meaningful probabilistic value of resistance cost reflecting a tolerable likelihood for resistance selection.

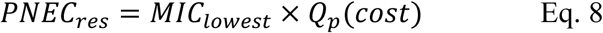

### Proposed methodology for PNEC_res_ estimation

#### Lowest known MICs

The database of MIC distributions hosted and maintained by EUCAST^27^ is nowadays the most extensive resource to obtain estimates of MIC_lowest_. In 2016, Bengtsson-Palme & Larsson^29^ proposed an algorithm yielding estimates of MIC_lowest_ for all antibiotics of clinical relevance, exploiting the EUCAST database. Briefly, for every combination of antibiotic and test species, a robust minimum estimate is extracted from the reported distribution of MIC values. The minimum MIC of the most sensitive species is considered a preliminary estimate of MIC_lowest_ for a particular antibiotic. Such preliminary estimates of MIC_lowest_ are obviously more representative if more distinct species were tested. Consequently, the preliminary estimates of MIC_lowest_ undergo a final adjustment to compensate for the expected positive bias in cases of low species coverage.

For this study, we reimplemented the algorithm of Bengtsson-Palme & Larsson^29^ in the R language^41^ for the sake of improved code transparency and modularization^42^. The preliminary estimates of MIC_lowest_ computed from a recent (2024) EUCAST dataset^27^ were identical to the estimates obtained by Bengtsson-Palme & Larsson in 2016^29^ for the majority of antibiotics (73 out of 106 cases; Fig. 4). Deviations hardly exceed factor four except for a few outliers explicitly labelled in Fig. 4. Overall, the comparison reveals a slight but significant trend towards lower MIC_lowest_ values (p = 0.0005, Wilcoxon signed-rank test) reflecting a better coverage of extremes by the grown MIC database (MIC_lowest_ declined for 25 out of 106 antibiotics, an increase was observed for 8 out of 106 only).

**Figure 4:**
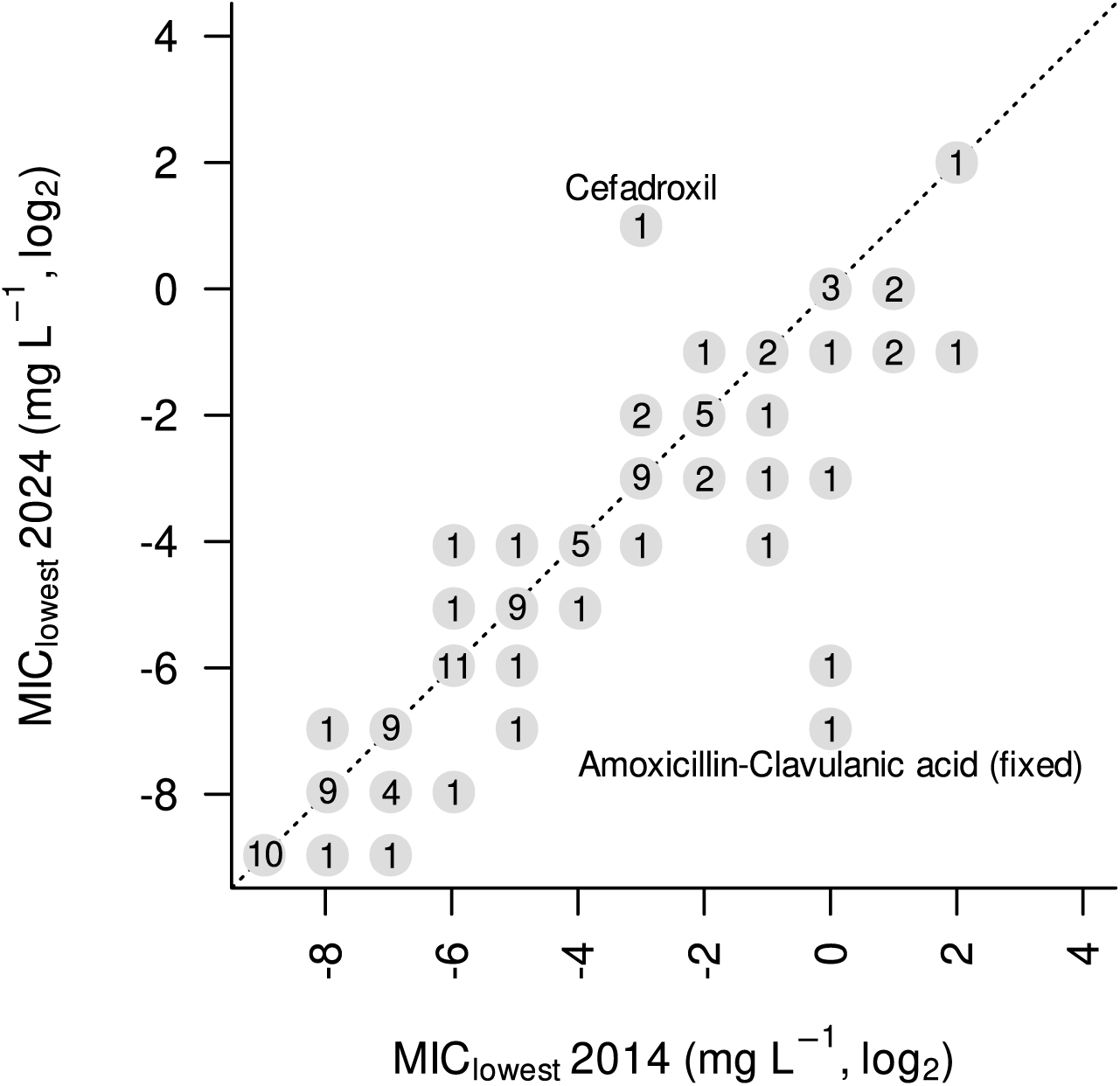
Lowest MICs computed from a 2024 snapshot of the EUCAST data in comparison to the lowest MICs reported by Bengtsson-Palme & Larsson in 2016^29^ prior to adjustment for species coverage. Integers represent the number of antibiotics where MIC_lowest_ experienced the respective shift (names are indicated for the two most extreme shifts only). Example: According to the 2014 data, eight antibiotics were associated with MIC_lowest_ = 2^-2^ mg/L. An update to the 2024 dataset resulted in a decline by one unit (new MIC_lowest_ = 2^-3^) for two of them, an increase to 2^-1^ was observed for a single antibiotic, and MIC_lowest_ did not change for the five remaining antibiotics. Regular distances between dots reflect the resolution of the EUCAST test scale.

Our reimplementation^42^ of the Bengtsson-Palme & Larsson algorithm^29^ differs slightly from the original regarding the compensation of low species coverage. While the original algorithm employs a discontinuous linear model to adjust the preliminary estimates of MIC_lowest_, the reimplementation adopts a continuous non-linear bias correction model (see Text S2 and Fig. S2).

#### Distribution of resistance cost

To the best of our knowledge, a curated and comprehensive database of antibiotic resistance costs does currently not exist. A previous compilation of resistance costs by Vogwill & MacLean^37^ provides an excellent starting point for such a database. We complemented the latter with information from additional recent scientific publications as well as our observations underlying Figure 3. The available data (220 records from 82 distinct studies) represent both plasmid-borne and chromosomal resistance elements, but taxonomic coverage in the literature is uneven with a strong bias towards selected pathogens, particularly *E. coli* (Fig. 5A).

**Figure 5:**
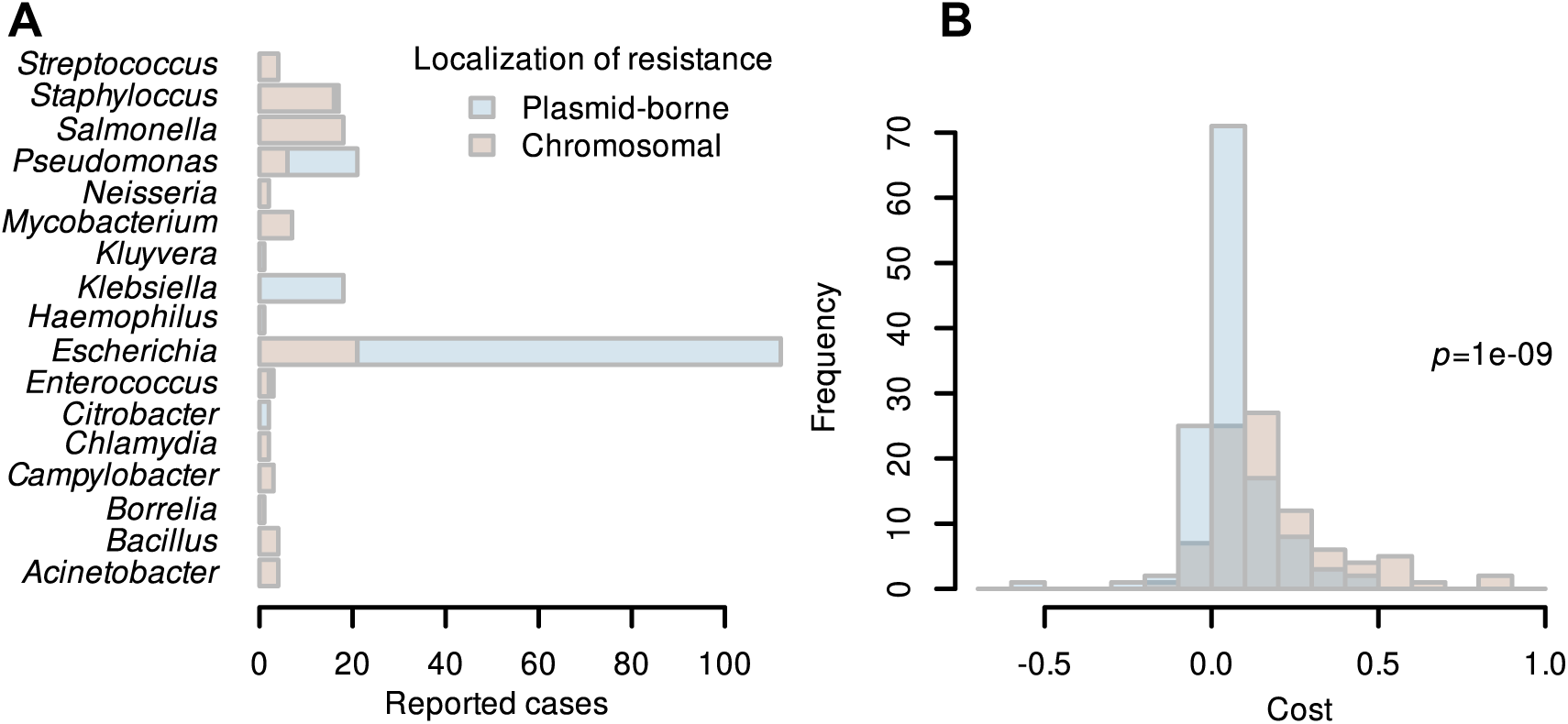
A) Available estimates of resistance costs for different bacterial genera and genetic localizations of the resistance. B) Fitness costs associated with antibiotic resistance reported in the scientific literature (n=194) and observed in own competition experiments (n=26, data from Fig. 3). Colour indicates whether the resistance mechanism is coded on chromosomal DNA or plasmids.

These resistance costs form a right-skewed distribution irrespective of whether resistance is encoded on a plasmid or chromosome. Chromosomal resistances exhibit a higher fitness cost when compared with plasmid-borne resistance (*p* < 10^-6^, Wilcoxon rank sum test), indicated by shifted histograms (Fig. 5B). This is according to ecological theory, as plasmid-encoded AMR can be horizontally transferred, allowing strains to acquire resistance without undergoing potentially detrimental mutations^43^. Moreover, plasmids frequently carry regulatory mechanisms that optimize gene expression, ensuring that ARGs are only activated when needed, thereby minimizing metabolic burden^44^. Additionally, plasmids can evolve compensatory adaptations over time, reducing their fitness cost, while chromosomal resistance mutations often disrupt essential cellular processes, leading to persistent growth disadvantages^39,40,43,44^.

About 10% of the reported costs in the literature are negative (Fig. 5B) indicating an apparent increase in the strain’s growth rate upon acquisition of resistance (Eq. 2). We propose that such negative cost estimates reflect either experimental uncertainty or an actual gain in fitness attributed to the acquisition of a plasmid. Plasmid-encoded resistance can sometimes provide a growth benefit even in the absence of selective pressure through the antibiotic due to additional genes that enhance bacterial fitness. Many resistance plasmids also carry genes involved in stress response, virulence, biofilm formation, or metabolic adaptation, which can improve bacterial survival under diverse conditions^43^. Additionally, bacteria rapidly acquire compensatory mutations that reduce or even reverse the fitness costs associated with plasmid carriage, sometimes leading to increased growth rates^45^. Furthermore, plasmids often encode toxin-antitoxin systems that stabilize their inheritance and can enhance persistence under stress^46^. These factors explain why plasmid-carrying bacteria may occasionally outcompete plasmid-free counterparts, even without direct antibiotic selection. Consequently, for these cases, avoiding positive selection through the regulation of environmental antibiotic concentrations is impossible, and the focus of regulatory intervention should be mainly on those cases with a positive cost of resistance.

When estimating the second factor of Eq. 8, we exclusively focused on reported costs of plasmid-borne resistances (blue shade in Fig. 5B). The rationale behind the plasmid-centric view was the much greater risk of instant and fast proliferation of resistance across multiple hosts enabled through horizontal gene transfer^47,48^ as compared to chromosomal mechanisms together with their slightly lower cost.

The focus on particularly "cheap" plasmid-borne resistance determinants necessarily triggers the question of whether MIC and cost data are independent such that a multiplication of two lower-bound estimates (Eq. 8) is justified. Given the limited set of genera for which resistance costs have been measured (Fig. 5A), the question cannot be answered exhaustively. It can be shown, however, that plasmid-borne resistance, which is typically low-cost (Fig. 5B), regularly occurs in bacterial genera whose wild-type strains exhibit the strongest known sensitivity to the respective antibiotics (Fig. 6).

**Figure 6:**
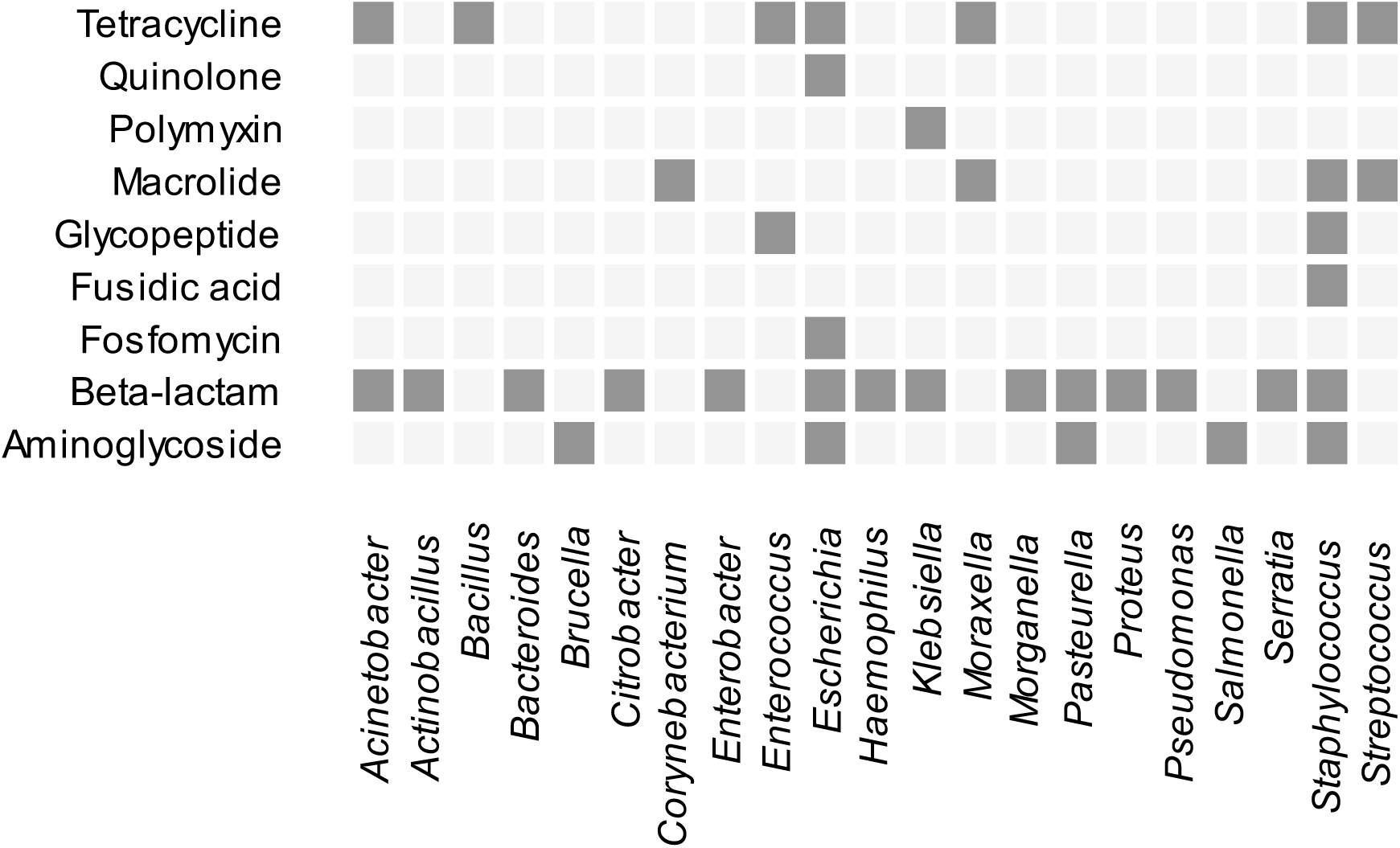
Co-occurrence of exceptional antibiotic sensitivity and plasmid-borne ARGs. Dark-shaded cells indicate that members of the respective genus are the most sensitive organisms concerning antibiotics of the corresponding class (inferred from EUCAST MIC data^27^). At the same time, isolates of that genus have been shown to carry plasmid-borne ARGs targeting the particular class of antibiotics. For this analysis, all plasmid sequences present in PLSDB^49–51^ were scanned for acquired ARGs covered by the resfinder database (version 2.1.1)^52^.

Based on this evidence that the multiplication of MIC_lowest_ and costs is reasonable, we aimed at identifying the probability distribution of actual resistance costs against the background of partly negative estimates. We hence approximated the distribution by a mixture model with two components: (1) an exponential distribution reflecting the actual cost of resistance and (2) a normal distribution centred around zero accounting for observed fitness alterations not strictly attributable to resistance (Fig. 7). The latter component pragmatically addresses both experimental uncertainties and alterations in fitness coinciding with resistance acquisition.

**Figure 7:**
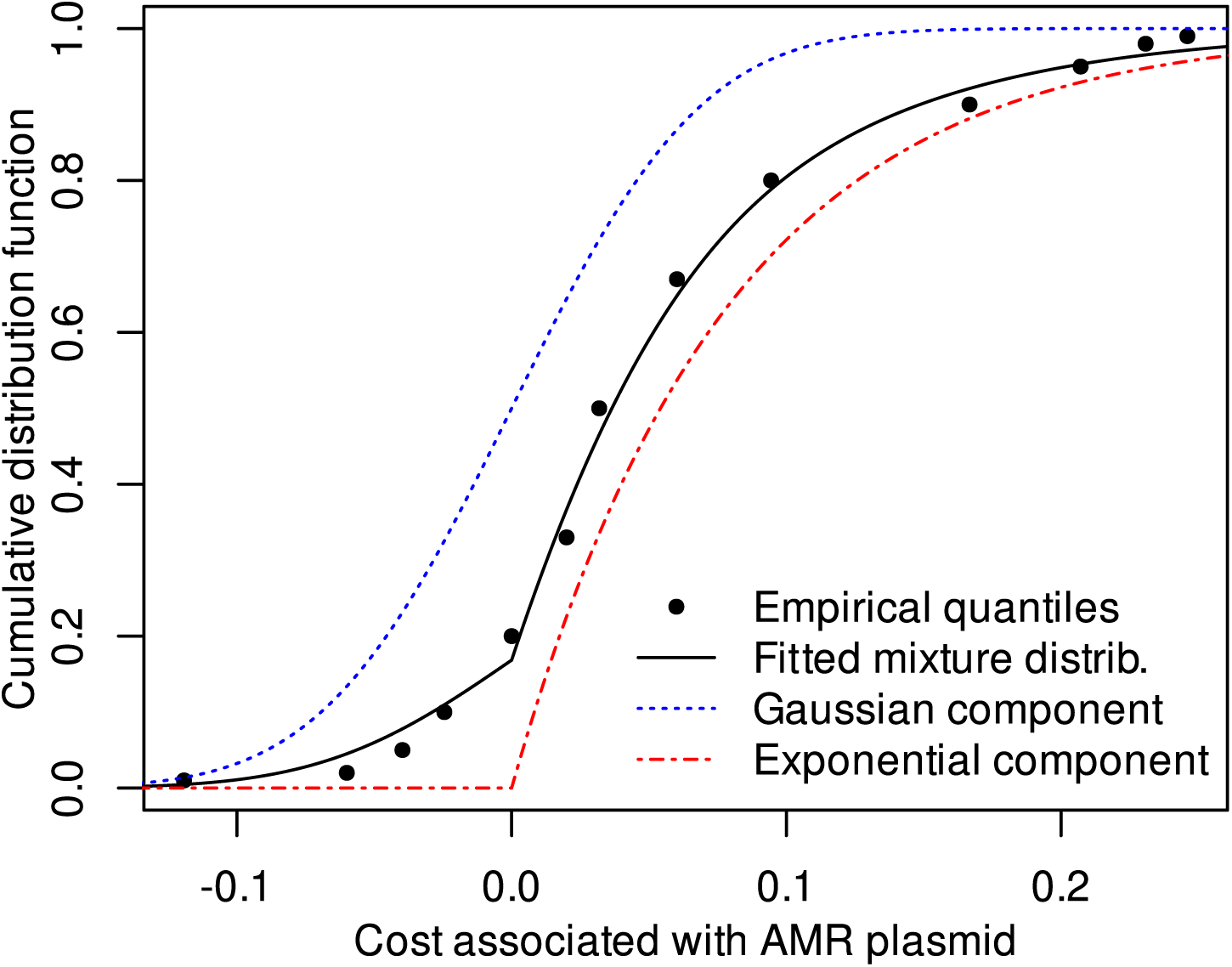
Approximation of the cost distribution observed in competition experiments between strains with a plasmid-borne resistance and its susceptible counterpart (n=122 after omission of 8 outliers with abs(cost) > 0.25). The empirical distribution of cost data (quantiles shown as dots) was approximated by a mixture distribution model (black line). The latter represents a weighted sum of two basic distribution models, termed components. The distribution of the actual resistance costs (strictly positive) is represented by the exponential component (red line). The zero-centred Gaussian component (blue) accounts for both noise in empirical data as well as alterations in fitness coinciding with resistance acquisition. Such apparent or actual alterations in fitness can be positive or negative. The three parameters of the mixture distribution (one for each component plus a weight parameter) were fitted by a conventional maximum likelihood approach.

By evaluating the exponential component of the mixture distribution reflecting the loss of fitness strictly attributable to resistance (Fig. 7), cost estimates can be obtained for any probability (Table 1) reflecting the intended level of protection. Based on the available data, the costs associated with a probability threshold of, e.g., 5% of resistance determinants still being selected for would be approximately 1/250. Hence, if a 5% probability threshold was considered acceptable from a risk management perspective, Eq. 8 would predict the PNEC_res_ to be about 1/250 of MIC_lowest_ (Table 1). Note that the determination of an acceptable probability threshold regarding the level of protection for regulatory purposes is beyond the scope of this manuscript, as it involves a risk-benefit assessment with the appropriate consideration of, among others, ethical and financial aspects as described elsewhere^53^.

**Table 1:** Association of plasmid-borne resistance costs with probabilities as inferred from the exponential component of the mixture distribution of Fig. 7. Costs are presented as a percentage for convenience (values according to Eq. 2 multiplied by 100).

| Probability | Cost (%) | 95% Confidence interval |
| --- | --- | --- |
| 0.01 | 0.079 | 0.063 - 0.1 |
| 0.02 | 0.16 | 0.13 - 0.2 |
| 0.05 | 0.4 | 0.32 - 0.52 |
| 0.1 | 0.82 | 0.66 - 1.1 |
| 0.25 | 2.2 | 1.8 - 2.9 |
| 0.5 | 5.4 | 4.3 - 7.0 |

#### Necessity of an ecotoxicological assessment factor

When deriving PNEC values, it is standard ecotoxicological practice to apply assessment factors (AFs) to account for uncertainty and variability when extrapolating from limited laboratory data to complex natural ecosystems^54^. In principle, the derivation of PNEC_res_ in Eq. 8 could similarly be extended by incorporating an AF to ensure that the resulting values are protective against selection across diverse environmental species and resistance determinants. Previous efforts - such as those by Bengtsson-Palme & Larsson^29^ and Rico et al.^31^ - have used an AF of 0.1, primarily to account for the discrepancy between MICs and MSCs.

In our approach, however, such an adjustment is unnecessary, as Eq. 8 already incorporates a conservative lower-bound estimate of the cost of resistance to convert MIC to MSC. Therefore, any additional AF would only need to address residual uncertainties in the environmental distributions of MICs and resistance costs. Given the breadth and depth of the underlying MIC and resistance cost datasets, we consider these estimates to be relatively robust. Furthermore, since Eq. 8 multiplies the lower-bound estimates of both parameters, the likelihood of underestimating the threshold for selection is further reduced. Based on these considerations, we propose using an AF of 1 when deriving PNEC_res_ from Eq. 8 - effectively omitting the AF - while emphasizing that this recommendation should be revisited during regulatory evaluation processes.

### Comparison with existing frameworks

The basic structure of Eq. 8 is identical to existing approaches for the estimation of PNEC_res_ addressing selection for AMR^29,31,32^ with novelty being limited to the adjustment applied to MIC_lowest_. Bengtsson-Palme & Larsson proposed PNEC_res_ = MIC_lowest_ / 10 ^29^. They denoted the number 1/10 a "flat assessment factor" to account for differences between MICs and MSCs without providing a specific justification based on empirical data or ecological theory. The very same assumption (MSC ∼ MIC / 10) was adopted by Rico et al.^31^ with reference to empirical data by Gullberg^8^ and Liu^9^. Given the probabilistic cost estimates of Table 1, it is clear that our study calls for a rethinking of the factor of 10 which has recently been adopted in a WHO guidance document on wastewater and solid waste management for manufacturing of antibiotics^55^ as well as by the AMR Industry Alliance for determining antibiotic thresholds for pharmaceutical manufacturing discharges released into surface waters^30^.

If, for example, the level of protection should cover 95% of possible resistance costs, a more justifiable estimate of PNEC_res_ would be MIC_lowest_ / 250, as the corresponding cost estimate where 5% of resistances are below the threshold is about 0.004 (0.4 %) (Table 1). If the presented framework is used to compute a PNEC_res_, the obtained value will potentially differ from the previously proposed values for two reasons: (1) an altered estimate of MIC_lowest_ and (2) the newly proposed conversion factor based on resistance cost. This can, for example, be illustrated with the case of Doxycycline. Based on the EUCAST data from 2014 evaluated by Bengtsson-Palme and Larsson in 2016, the primary estimate of MIC_lowest_ was 32 µg/L, which was adjusted to 16 µg/L in consideration of the limited number of tested species (29 at that time with highest sensitivity seen in *Campylobacter jejuni*, *Staphylococcus aureus*, and *Streptococcus pyogenes*). The application of the assessment factor of 10 followed by rounding resulted in a PNEC_res_ of 2 µg/L. In the 2024 version of EUCAST, the number of species tested for Doxycycline resistance has grown to 55, accompanied by a reduction of MIC_lowest_ to 8 µg/L, owing to the susceptibility of *Bacillus anthracis*. Considering the cost-based conversion factor of 250 leads to an updated PNEC_res_ of 0.032 µg/L.

## Conclusion

### Scenarios of selection and applicability of the proposed framework

Upon release into the environment, antibiotics potentially select for contrasting bacterial features that must be clearly distinguished: (1) general non-susceptibility, (2) immobile chromosomal resistance, and (3) mobile resistance.

General non-susceptibility addresses all cases where bacteria of a certain phylogenetic branch lack the target of the antibiotic in question. While the respective bacterial genera can profit from antibiotic exposure indirectly via competition, there is no immediate risk associated with their selection as no inheritable ARGs are involved. Bacterial genera carrying immobile chromosomal ARG can likewise profit from antibiotic exposure and undergo an enrichment within communities over time. Besides the increased abundance of vertically inheritable ARGs, the enrichment is possibly linked to an elevated but still rare chance of stochastic mobilization^56^, making it more likely for the ARG to emerge in human and veterinary pathogens in the long run. Finally, antibiotics exposure selects for mobile ARGs, especially those harboured by plasmids, many of which can be acquired by a broad spectrum of bacterial hosts^48,57–59^. The presence of plasmid-borne resistance in environmental settings typically reflects pollution scenarios where both antibiotics and pathogens with a history of exposure are released simultaneously through, e.g., wastewater disposal^60^ or organic fertilization^61^. Crucially, newly introduced plasmid-borne ARGs can be transferred laterally to species that constitute typical members of genuine environmental microbiomes^48^. As a consequence of such lateral transfer, critical ARGs can persist and multiply in water and soil over extended periods of time and undergo particularly efficient proliferation under selective conditions.

On short- and medium-term time scales, the selection of plasmid-borne mobile ARGs is considered the scenario of primary relevance as it allows for the fastest proliferation and often triggers the simultaneous spread of multiple health-critical traits. Consequently, the factor to convert MIC_lowest_ into PNEC_res_ (Table 1) was chosen to primarily address this "plasmid scenario". Nevertheless, the principal framework expressed by Eq. 6 and Eq. 7 would be equally applicable to cases of immobile chromosomal resistances if the distinct distribution of costs (Fig. 5B) was taken into account.

With regard to estimates of MIC_lowest_, we still have to state an obvious underrepresentation of truly benign environmental bacteria in the underlying databases. Given the lack of immediate health relevance and considering the significant proportion of non-culturable genera, this underrepresentation is expected to persist. However, we are not aware of evidence for systematically lower MICs in benign environmental bacteria compared to facultative or obligate pathogens covered by large-scale susceptibility testing.

### Towards higher accuracy in estimated PNEC_res_

At the time of this writing, the amount and quality of information regarding the two factors of Eq. 8, namely the lowest MIC and the cost of resistance, is clearly imbalanced. While MIC data are collected and analysed routinely following strict standards, this is currently not the case for resistance costs. We believe that the accuracy of probabilistic information on resistance costs (Fig. 7, Table 1) could profit substantially from wider coverage of test organisms and genetic elements as well as standardized culturing protocols reflecting environmental scenarios. An aspect deserving particular attention is the selection of pairs of test strains. Apart from the resistance element under consideration, the strains must obviously be isogenic to allow for reasonable cost estimates. While plasmid transfer by conjugation or transformation is a straightforward means to create such strain pairs, plasmid stability in the recipient needs to be checked carefully. Ideally, the newly created resistant strain would be allowed to undergo adaptation to the plasmid over many generations to account at least for the very quick and highly replicable cases of resistance cost amelioration^62^.

Given the absence of high-quality information on the shape of dose-response curves for the vast majority of combinations of antibiotics and strain, the linear model (Eq. 1) was chosen as the consensus model as it mediates between the different curve shapes. Besides that, the linear model does not introduce any additional parameters in Eq. 6 or 7, beyond MIC and resistance cost. Nevertheless, if a non-linear consensus dose-response model provides a better fit to empirical data of a particular antibiotic or class of antibiotics, the respective model should be adopted to avoid a possible bias in the estimates of PNEC_res_. Note that the bias arising from the application of the linear model to cases where the actual dose-response curve is nonlinear can be positive or negative (see Fig. S1). Consequently, uncertainty introduced by the choice of a particular dose-response relationship can never be cured by means of a universal assessment factor but requires treatment at the level of individual antibiotics or antibiotic classes as outlined in the supplementary material (Text S1 and Eq. S1a, S1b).

### Implementation and re-evaluation

At present, the cost estimates provided in Table 1 and PNEC_res_ derived thereof (Eq. 8) must consequently be treated as preliminary. Nevertheless, we do propose their immediate use in environmental regulation after deciding on appropriate probability thresholds reflecting the tolerable risks. This recommendation of immediate use is built upon the notion that the current practice (PNEC_res_ = MIC_lowest_ / 10) conflicts with precautionary principles very obviously. The adoption of the presented, cost-based approach to derive PNEC_res_ would not only help to protect against resistance selection in the environment but could also foster trust in maximum permissible concentrations when the latter are derived from a fundamental, easily communicable ecological concept.

Finally, it must be acknowledged that the statistical distributions of both MICs and resistance costs are dynamic features. On the one hand, moderate changes in distribution parameters may occur in response to continued data collection. On the other hand, gradual changes in antibiotic susceptibility and resistance costs over time reflect the continuous evolution of bacterial genomes and mobile genetic elements. Consequently, a periodic revaluation of PNEC_res_ followed by an adjustment of regulations is inevitable.

## Online Methods

### Experiments

#### Identification of minimum inhibitory concentrations (MIC)

MIC determination was performed following the EUCAST guidelines for broth microdilution assays^63^. Briefly, wild-type bacteria were cultured in Müller-Hinton (MH) broth overnight at 37 °C under agitation (120 rpm), then diluted to a standard concentration of 5 x 10⁵ CFU/ml for consistency. Fresh antibiotic stock solutions were prepared according to manufacturer or CLSI guidelines. Serial two-fold dilutions were made in MH broth, with concentrations based on EUCAST breakpoints for the specific antibiotic and bacterial species. The diluted bacterial suspension was added to a microtiter plate containing the antibiotic dilutions, with four replicates for each antibiotic concentration. Plates were incubated at 35 ± 2 °C for 18 h to allow for bacterial growth. After incubation, the MIC was identified as the lowest antibiotic concentration that completely inhibited visible bacterial growth based on spectrophotometric optical density measurements at a wavelength of 600 nm (OD_600_) (Synergy H1, BioTek Instruments, Inc., Winooski, VT, USA). Controls (growth control, sterility control) and reference strains with known MIC values were included. MIC determination was also performed at 30 °C, with no significant differences in MIC values observed compared to those determined at the EUCAST-suggested temperature.

#### Preliminary estimates of the minimum selective concentration (MSC_approx_)

Competition experiments targeted at identifying the MSC require antibiotic concentrations to be varied around that critical concentration where the resistant and susceptible strain exhibit equal fitness. To confine the range of concentrations to be actually tested, we first calculated an approximation (MSC_approx_) making use of Eq. 7. Besides the respective MICs (see above), Eq. 7 requires the cost of resistance as an input. An approximate estimate of the cost was obtained by comparing the two strains’ intrinsic growth rates at zero antibiotic exposure (Eq. 2). For the latter purpose, growth curves of both strains at 30 °C were recorded in a microplate reader (Synergy H1) by measuring the OD_600_ (30 replicates per strain). Growth rate constants were obtained as the maximum slope of a linear model fitted to log-scaled OD_600_ values recorded during the exponential phase.

#### Identification of minimum selective concentrations (MSC) by competition

Empirical MSCs were determined through competition experiments using pairs of isogenic bacterial strains that differed in their resistance phenotype only (Table 2). Both bacterial strains (resistant and susceptible) were cultured individually overnight in MH broth at 30 °C under constant agitation (120 rpm). The cultures were then diluted 1:10⁶ in fresh MH broth to standardize the initial bacterial density. The diluted bacterial suspensions were mixed at a 1:1 ratio and incubated without antibiotics (control) as well as in broth amended with antibiotics (concentrations varied around MSC_approx_ in increments of factor 5). The co-cultures were left under constant agitation (120 rpm) at 30°C for 24 hours. Significant oxygen depletion was prevented through culturing in glass vials of 100 mL capacity filled with 20 mL of culture. All experiments were performed in six replicates. The initial densities of the two competing strains were quantified by individual plating before mixing on LB agar. After 24 hours of competition, bacterial counts for each strain were either obtained by plating on LB agar (if strains could be distinguished by fluorescence) or by plating on MH agar with and without antibiotic (for strain pairs without fluorescence markers). For each of the tested concentrations (c), the fitness ratio (fr) was computed from the abundances of the resistant and susceptible strain (a_res_, a_sus_) (Eq. 9), with the empirical MSC determined as the concentration where fr = 1 after fitting the obtained fr(c) data by a monotone Hermite spline^64^.

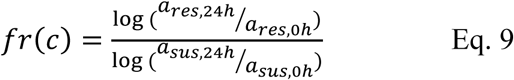

**Table 2:** Competition assays performed with strain pairs of Bacillus subtilis, Escherichia coli, and Pseudomonas putida to verify the predictions by Eq. 7.

| Bacterial strain | Resistance vector | Resistance gene | Antibiotic |
| --- | --- | --- | --- |
| *B. subtilis* W168 | Chromosomal <sup>a</sup> | *aph(3')-III* | Kanamycin |
| *B. subtilis* W168 | Chromosomal <sup>a</sup> | *erm(C)* | Clarithromycin |
| *B. subtilis* W168 | Chromosomal <sup>a</sup> | *erm(C)* | Tylosin |
| *E. coli* K12 | pB10 plasmid <sup>65</sup> | *aph(3'')-Ib, aph(6)-Id* | Streptomycin |
| *E. coli* K12 | pB10 plasmid <sup>65</sup> | *tetA* | Tetracycline |
| *E. coli* K12 | pB10 plasmid <sup>65</sup> | *bla<sub>OXA-2</sub>* | Amoxicillin |
| *E. coli* K12 | pB10 plasmid <sup>65</sup> | *tetA* | Chlortetracycline |
| *E. coli* K12 | pB10 plasmid <sup>65</sup> | *tetA* | Oxytetracycline |
| *E. coli* K12 | pB10 plasmid <sup>65</sup> | *tetA* | Doxycycline |
| *E. coli* K12 | RP4 plasmid <sup>66</sup> | *tetA* | Tetracycline |
| *E. coli* K12 | RP4 plasmid <sup>66</sup> | *aph(3')* | Kanamycin |
| *E. coli* K12 | MDR plasmid <sup>b</sup> | not identified | Trimethoprim |
| *E. coli* K12 | MDR plasmid <sup>b</sup> | not identified | Piperacillin |
| *E. coli* K12 | MDR plasmid <sup>b</sup> | not identified | Amoxicillin |
| *E. coli* K12 | MDR plasmid <sup>b</sup> | not identified | Tetracycline |
| *E. coli* K12 | Chromosomal <sup>c</sup> | *catA1* | Chloramphenicol |
| *E. coli* K12 | Chromosomal <sup>c</sup> | *aacC1* | Gentamicin |
| *P. putida* KT 2440 | pB10 plasmid <sup>65</sup> | *tetA* | Tetracycline |
| *P. putida* KT 2440 | pB10 plasmid <sup>65</sup> | *aph(3'')-Ib, aph(6)-Id* | Streptomycin |
| *P. putida* KT 2440 | pB10 plasmid <sup>65</sup> | *bla<sub>OXA-2</sub>* | Amoxicillin |
| *P. putida* KT 2440 | pB10 plasmid <sup>65</sup> | *tetA* | Chlortetracycline |
| *P. putida* KT 2440 | pB10 plasmid <sup>65</sup> | *tetA* | Oxytetracycline |
| *P. putida* KT 2440 | pB10 plasmid <sup>65</sup> | *tetA* | Doxycycline |
| *P. putida* KT 2440 | RP4 plasmid <sup>66</sup> | *tetA* | Tetracycline |
| *P. putida* KT 2440 | RP4 plasmid <sup>66</sup> | *aph(3')* | Kanamycin |

a: Strains created by Denis Iliasov, TU Dresden, Institute of Microbiology. b: Unidentified multidrug resistance plasmid captured from municipal wastewater by Serena Caucci, former member of the TU Dresden Institute of Hydrobiology. c: Strains chromosomally engineered using the pMRE-Tn5-145 delivery plasmid<sup>67</sup>.

### Data analysis

Statistical data analysis and visualization were performed using "R" 4.4.2 (www.r-project.org). The "bbmle" package^68^ (version 1.0.25.1) was employed for fitting distribution models by maximum likelihood. 95% confidence intervals of the resistance costs associated with particular probability levels (Table 1) were constructed by ordinary bootstrapping. The obtained bootstrap estimates were validated against alternative estimates inferred from the 95% CI of the fitted parameter of the exponential distribution generated by the profile likelihood method. The analysed collection of empirical resistance costs reported in the scientific literature is made available as part of the public repository https://github.com/enviresist/PNEC_res. The attached metadata include the bacterial species, the localization of the resistance determinant (plasmid or chromosome), the studied antibiotic(s), and the original data source (DOI).

The presence of plasmid-encoded antibiotic resistance in highly sensitive bacterial genera tested by EUCAST (Fig. 6) was studied by screening the plasmid sequence database "PLSDB"^51^ (https://ccb-microbe.cs.uni-saarland.de/plsdb2025, version from November 2023) for acquired resistance genes registered in the "resfinder" database^52^ (http://genepi.food.dtu.dk/resfinder, version 2.1.1). Sequence alignments were performed with NCBI’s "blastn" (version 2.9.0).

### Availability of data and source code

Snapshots of the processed EUCAST MIC data from 2014 and 2024, the database of resistance costs built from published values, and the R source code to derive PNEC_res_ is made available via the GitHub repository https://github.com/enviresist/PNEC_res^42^ managed by the first author. The computed values of PNEC_res_ can also be looked up via the interactive web interface accessible at https://enviresist.github.io/PNEC_res.

## Acknowledgements

The authors thank Mitja Remus-Emsermann, Denis Iliasov and Serena Caucci for access to, design and original isolation of strains used in this study. We further thank Juliane Isler, Steffen Kunze, Christiane Zschornack, and Melanie Tannert for technical lab support.

## Author contributions

**DKn**: Conceptualization, Methodology, Software, Validation, Formal analysis, Investigation, Resources, Data Curation, Writing - Original Draft, Writing - Review & Editing, Visualization, Supervision, Project administration, Funding acquisition; **MCB**: Conceptualization, Methodology, Validation, Formal analysis, Investigation, Resources, Data Curation, Writing - Original Draft, Writing - Review & Editing; **DKo**: Methodology, Investigation, Writing - Review & Editing; **VW**: Investigation; **PS**: Conceptualization, Writing - Review & Editing, Project administration, Funding acquisition; **KWS**: Conceptualization, Writing - Review & Editing, Project administration, Funding acquisition; **JS**: Conceptualization, Writing - Review & Editing, Project administration, Funding acquisition; **DJ**: Conceptualization, Writing - Review & Editing, Project administration, Funding acquisition; **TUB**: Conceptualization, Writing - Review & Editing, Project administration, Funding acquisition; **UK**: Conceptualization, Methodology, Software, Validation, Formal analysis, Investigation, Resources, Data Curation, Writing - Original Draft, Writing - Review & Editing, Visualization, Supervision, Project administration, Funding acquisition

