## Supplemental Information for "Suppressing selection for antibiotic resistance in the environment: A transparent, ecology-based approach to predicted no-effect concentrations"

***Text S1: Variation of dose-response curves between antibiotics and impact of the model choice on MSC estimates***

The proposed equations to calculate the MSC from the MIC (Eq. 6 & 7) are conditioned on a linear relation between growth rate and antibiotic concentration (Eq. 1). Actual dose-response curves are known to differ between classes of antibiotics (Fig. S1) and substantial inter-species variance is to be expected. Against this background, the linear dose-response model was chosen as a minimal consensus. As a generalization of Eq. 1, the model can be expressed by Eq. S1a with the exponent ß fixed at a value of 1.

$$\mu(c) = \mu^0 \times \left[1 - \left({}^{c}/_{MIC}\right)^{\beta}\right] \qquad for\ c < MIC \qquad \text{Eq. S1a}$$

On the one hand, the linear dose-response model (ß = 1) is the most parsimonious one in terms of parameters. On the other hand, the linear shape represents an intermediate case falling between the contrasting extremes seen for different antibiotics. For Mecillinam, for example, a proper fit of Eq. S1a requires ß > 1 whereas the dose-response curve for Trimethoprim indicates ß < 1 (Fig. S1). According to Fig. S1, adoption of the linear model is appropriate for some antibiotics, but it can lead to either over- or underestimation of MSCs for other antibiotics. Since the respective bias can be both positive or negative, the partial deficiency of the linear model could only be cured at the level of particular antibiotics. Consequently, whenever representative information on the shape of dose-response curves is available, we recommend substituting the linear model with the respective non-linear alternative. The simplest option to account for non-linear dose-response curves is to adopt Eq. S1a without constraints on the single additional parameter (ß) and to fit its value to observations as done in Fig. S1. Ideally, observations would cover a diverse set of bacterial strains. Once the value of ß is known, the MSC can be calculated from Eq. S1b replacing Eq. 6. The derivation of Eq. S1b is conceptually identical to the derivation of Eq. 6. For the special case of ß = 1, Eq. S1b obviously collapses to Eq. 6.

$$ {}^{MSC}/_{MIC_{sus}} = \left[\frac{cost}{1 - {}^{1}/_{f^{\beta}} \times (1 - cost)}\right]^{1/\beta} \qquad \text{Eq. S1b} $$

***Text S2: Adjustment of preliminary estimates of MIClowest for species coverage***

Bengtsson-Palme & Larsson (2016)[29] adjusted the preliminary estimate of $MIC_{lowest}$ used Eq. S2a to account for uncertainty arising from a limited number of species (n) covered by the EUCAST dataset with f = n / 41 if n < 40 and f = 1 otherwise.

$$MIC_{lowest} = MIC_{lowest,prelim} \times f \qquad \text{Eq. S2a}$$

The empirical parameters of this discontinuous linear model (40, 41) were found by randomly subsampling the original dataset to mimic situations where MIC data are available for a limited number of test organisms only. The update of the EUCAST MIC database to the 2024 version called for a re-estimation of the empirical parameters. In that context, we decided to substitute the discontinuous linear model (Eq. S2a) by a continuous version (Eq. S2b)

$$MIC_{lowest} = {MIC_{lowest,prelim}}^{a} \Big/ \left(b + {MIC_{lowest,prelim}}^{a}\right) \qquad \text{Eq. S2b}$$

with the free parameters a and b. Compared to Eq. S2a the structure of Eq. S2b allows for a better approximation of the data on the one hand but it is also simpler to fit with standard approaches like, e.g., R's built-in "nls" function. For the 2024 EUCAST dataset, a=1.367 and b=57.26 provided the best fit to the resampling data (Fig. S2).

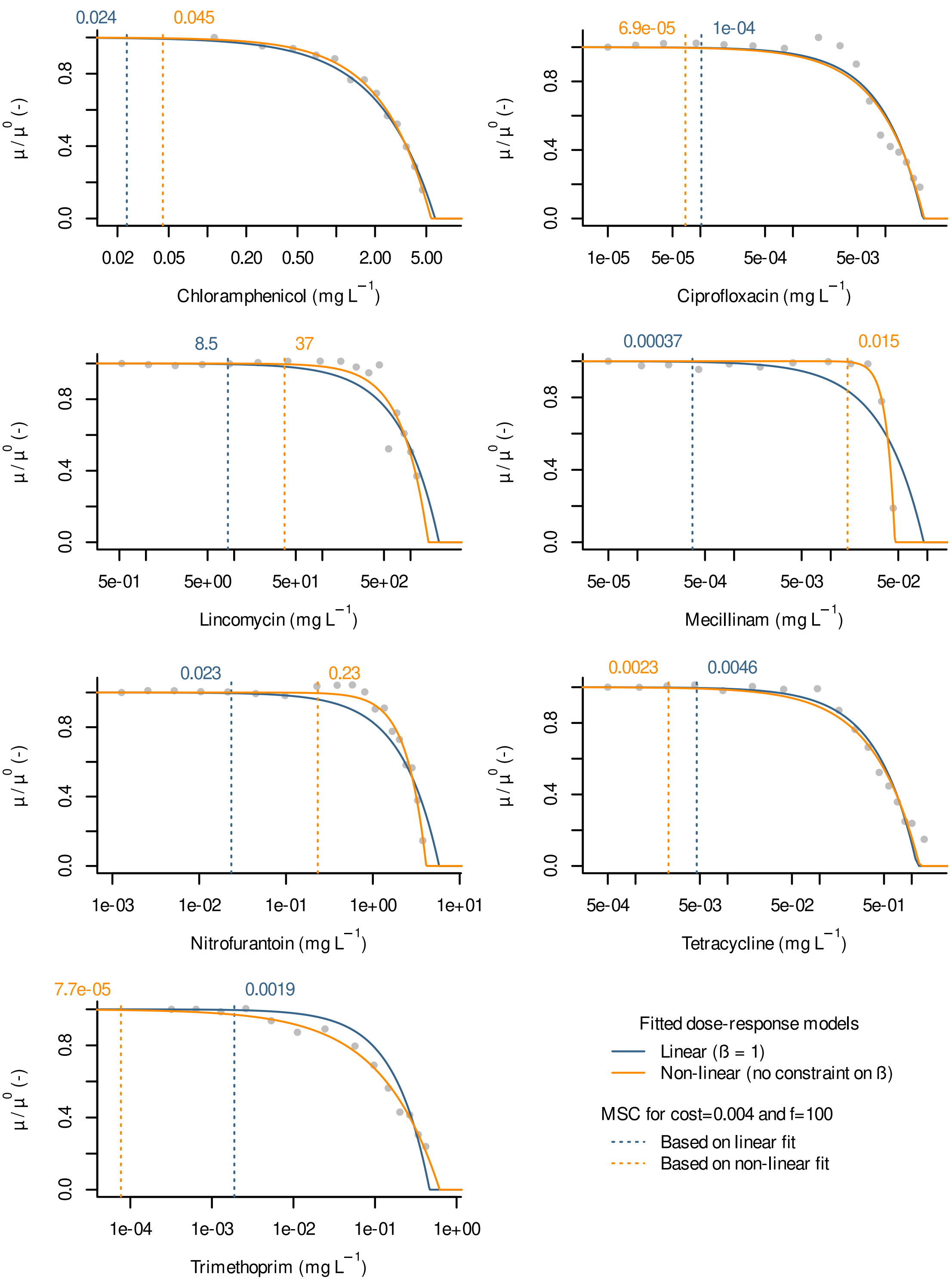


***Figure S1: Linear and non-linear dose response models (Eq. S1a) fitted to the E. coli dataset attached to Figure 1A of Angermayr et al., 2022[36]. Numbers above individual plots denote the estimated MSCs for a case of high-level resistance ($MIC_{res}$ = 100 * $MIC_{sus}$) and an assumed fixed cost of resistance of 0.004 representing the approximate 5% quantile of the distribution of plasmid-borne resistance costs according to Table 2 (calculated using Eq. S1b). Font colours correspond to the respective graphs.***

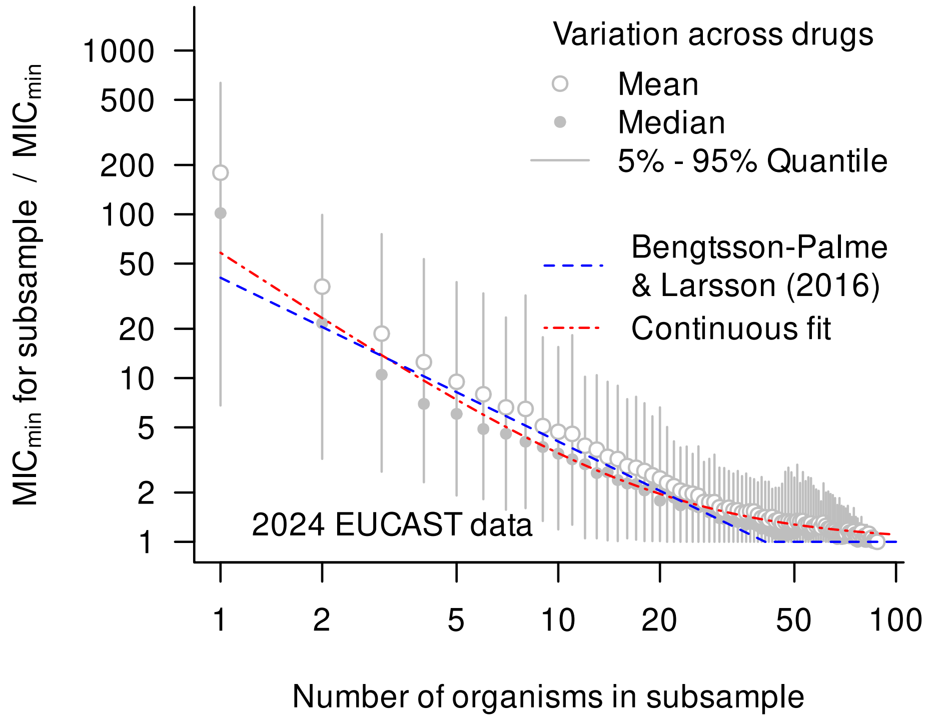


***Figure S2: Quotient of the minimum MIC of a subsample and the minimum MIC of the actual full sample (y axis) in relation to subsample size (x axis). Here, subsample size is identical to the number of distinct organisms (species). As the number of distinct organisms declines, the quotient increases systematically, indicating the need for a stronger downward adjustment of $MIC_{lowest}$. As compared to the original adjustment approach by Bengtsson-Palme and Larsson[29] (blue graph), the continuous model (red graph) results in a somewhat stronger correction in the extreme case of n=1 and it accounts for residual uncertainty at very high numbers of tested organisms in accordance with general probabilistic concepts.***
